# Regulation of arousal via on-line neurofeedback improves human performance in a demanding sensory-motor task

**DOI:** 10.1101/428755

**Authors:** J. Faller, J. Cummings, S. Saproo, P. Sajda

## Abstract

Our state of arousal can significantly affect our ability to make optimal decisions, judgments, and actions in real-world dynamic environments. The Yerkes-Dodson law, which posits an inverse-U relationship between arousal and task performance, suggests that there is a state of arousal that is optimal for behavioral performance in a given task. Here we show that we can use on-line neurofeedback to shift an individual’s arousal toward this optimal state. Specifically, we use a brain computer interface (BCI) that uses information in the electroencephalogram (EEG) to generate a neurofeedback signal that dynamically adjusts an individual’s arousal state when they are engaged in a boundary avoidance task (BAT). The BAT is a demanding sensory-motor task paradigm that we implement as an aerial navigation task in virtual reality (VR), and which creates cognitive conditions that escalate arousal and quickly results in task failure — e.g. missing or crashing into the boundary. We demonstrate that task performance, measured as time and distance over which the subject can navigate before failure, is significantly increased when veridical neurofeedback is provided. Simultaneous measurements of pupil dilation and heart rate variability show that the neurofeedback indeed reduces arousal. Our work is the first demonstration of a BCI system that uses on-line neurofeedback to shift arousal state and increase task performance in accordance with the Yerkes-Dodson law.

## Introduction

Why does walking across a brand new carpet with a full cup of coffee in one hand seem such a stressful and difficult task? If the cup is filled with water instead of coffee, and/or if the carpet is old and decrepit why does the task seem less daunting and less likely to result in a spill? The same can be said of the act of walking across a balance beam, where the difference in our performance — e.g. our speed across the beam and the likelihood of a fall — is dramatically lower if the beam sits six inches off the ground compared to when it is sixty feet up. Aphoristically, why do “high stakes” lead to “grave mistakes”?

One possible explanation invokes the deleterious impact of loss aversion on optimal cognitive control. Cognitive control typically refers to a set of cortical processes and neuromodulatory functions that configures cognition for optimal performance at a specific task (1, 2). When we are performing a high-consequence task, with performance boundaries that are critical — a spill of coffee out of the side of the cup that leads to spousal rebuke or the slip of the foot off the edge of the balance beam that leads to grave injury — arousal levels can increase sharply and cognitive control can be drastically diminished.

A highly specialized scenario that represents an extreme case of putting a high demand on sensory-motor cognition is related to an aviation phenomenon known as “pilot induced oscillation”, or PIO. PIOs are defined as unstable short-period oscillations in the motion of an aircraft, manifested by the pilot’s own control input. Spontaneous short-period oscillations are normal, but they can be catastrophic if the pilot over-compensates for small control errors in a way that increases the amplitude of these oscillations. PIOs have been simulated using a “boundary avoidance task” (BAT) paradigm (3–5). The BAT paradigm is thought to gradually increase a pilot’s cognitive workload, arousal, and task engagement, until cognitive control processes are overwhelmed and there is a catastrophic control failure, often resulting in a crash.

We recently conducted an investigation of PIOs using a naturalistic BAT paradigm where no feedback was provided (6, i.e. “open-loop”). We identified electroencephalographic (EEG) signatures that discriminated task difficulty level within a trial and showed that these signatures were predictive of an upcoming PIO event. The signatures were identified in a number of EEG frequency bands and spatial topographies, including frontocentral theta activity (4 to 7 Hz), occipital alpha activity (8 to 15 Hz),and posterior and temporal gamma band activity (32 to 55 Hz). The spatial and spectral pattern of the theta activity is consistent with engagement of the anterior cingulate cortex (ACC), a hub for cognitive control (7, 8). Occipital alpha, on the other hand has been tied to both arousal and visual selective attention (9, 10). Activity in the gamma band had topologies suggestive of muscle tension in the neck and head that, though not strictly cortical in origin, was informative of the upcoming PIO event. Using a linear decoder to combine this EEG information across frequency bands yielded the most robust predictor of an upcoming PIO (6). In addition these open-loop experiments showed correlations between subject’s pupil diameter and task difficulty, specifically increased pupil dilation as the task became more difficult and there was an increased likelihood of a PIO. This suggested that the subject’s state of arousal was changing during the task and there was a correlation between this state and the likelihood of task failure. In general, these open-loop observations are in-line with the well-known Yerkes-Dodson relationship (11, 12) between arousal and task performance (Fig. 1A).

**Fig. 1.**
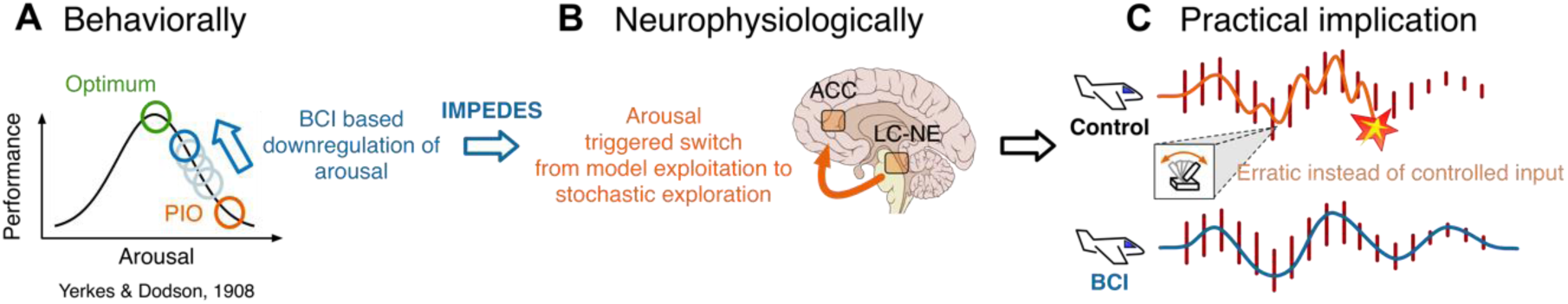
Assumptions underlying our hypothesis: (A) If performance decrease during pilot induced oscillations (PIO) is governed by the Yerkes-Dodson law, then down-regulation of arousal should improve performance. (B) The locus-coeruleus norepinephrine (LC-NE) system is believed to trigger a switch away from model exploitation to stochastic exploration, and thus hypothetically cause PIO, if arousal exceeds a threshold while anterior cingulate cortex is in a state reflective of low model performance (14). Lowering arousal should impede this switch to stochastic exploration and thus lower PIO propensity. (C) Subjects typically fail along the way in sufficiently difficult boundary avoidance tasks, but impeding PIOs by down-regulating arousal would hypothetically postpone failure and thus improve task performance.

In this paper we investigated whether the performance of subjects in a BAT could be improved using closed-loop neurofeedback that leverages these previously identified EEG signatures predictive of PIOs (Fig. 1). Here we define neurofeedback broadly, using signals decoded in the EEG bands (0.5 to 55 Hz, delta, theta, alpha, beta and gamma bands) that track task-dependent arousal state. As we observed in the open-loop experiments (6) the signatures we find and use for neurofeedback include sources in the central nervous system (CNS) as well peripheral nervous system (PNS) activity that is picked up in the EEG. We assessed improvement in performance by whether subjects could “fly” longer in difficult conditions (narrow boundaries) when veridical neurofeedback is provided relative to sham feedback or silence (Fig. 1C). The neurofeedback is based on information that we decode from the EEG in real-time via brain-computer interface techniques (BCI, (13)) and that we provide to subjects throughout the BAT. Specifically, feedback is given via headphones in the form of a loudness modulated low-rate (60 beats per minute) synthetic heartbeat. The loudness of the auditory feedback is inversely related to the tracked EEG signatures of task difficulty — i.e. louder feedback is provided when the level of inferred task-dependent arousal is high. Our hypothesis was that subjects would entrain to the low-rate heart beat when its loudness was increased during difficult moments in the trial. This would cause a reduction in arousal, which would shift performance on the task, in line with the Yerkes-Dodson relationship (Fig. 1 A and B). Indeed, our results show that subjects show a significant improvement in their task performance when using the closed-loop neurofeedback compared to when no feedback or sham feedback are provided. Furthermore, analysis of pupillometry and heart rate variability (HRV), neither of which is used to construct the neurofeedback signal, supports the hypothesis that the feedback is impacting performance by reducing arousal levels, consistent with models of cognitive control and the Yerkes-Dodson relationship (11, 12).

## Results

### Flight paradigm

Twenty healthy adults performed a BAT in a virtual reality (VR) environment, where they navigated a plane through courses of rectangular red waypoints (“rings”). Flight attempts (i.e. trials) were alternately performed in an easy and hard course (Fig. 2A, top left). Every course was a maximum of 90 seconds long, but ended abruptly whenever the pilot missed a ring. The size of the rings decreased every 30 seconds, thus increasing task difficulty. One of three feedback conditions (BCI, sham or silence) was randomly assigned for every new flight attempt: In the main condition of interest *BCI*, subjects heard audio of a low-rate synthetic heartbeat that was continuously modulated in loudness as a function of the level of inferred task-dependent arousal, as decoded from the electroencephalogram (EEG). The higher the level of inferred task-dependent arousal, the louder the feedback and vice versa. In the first control condition *silence*, no audio was presented. For the second control condition *sham*, the decoder-output was linearly combined in equal parts with random sham signal (see Section Feedback conditions). The linear decoder had before been trained based on spectral features of EEG collected during 10 minutes of flight attempts in the easy course at the beginning of the main experimental session (see Fig. 2B). Specifically, the decoder was trained to discriminate sections of EEG around large boundaries versus sections around medium/small boundaries. This was our proxy for EEG-derived arousal state that we hypothesized would couple to task performance. Subjects were kept blind with regard to the purpose of the study and the existence of the sham condition. The key instructions were as follows: “Consider missing a ring the equivalent of crashing a plane” and “Whenever you hear heartbeat audio, please try to assume a mental state where the audio becomes and stays as low in volume as possible” (see Methods for further details).

**Fig. 2.**
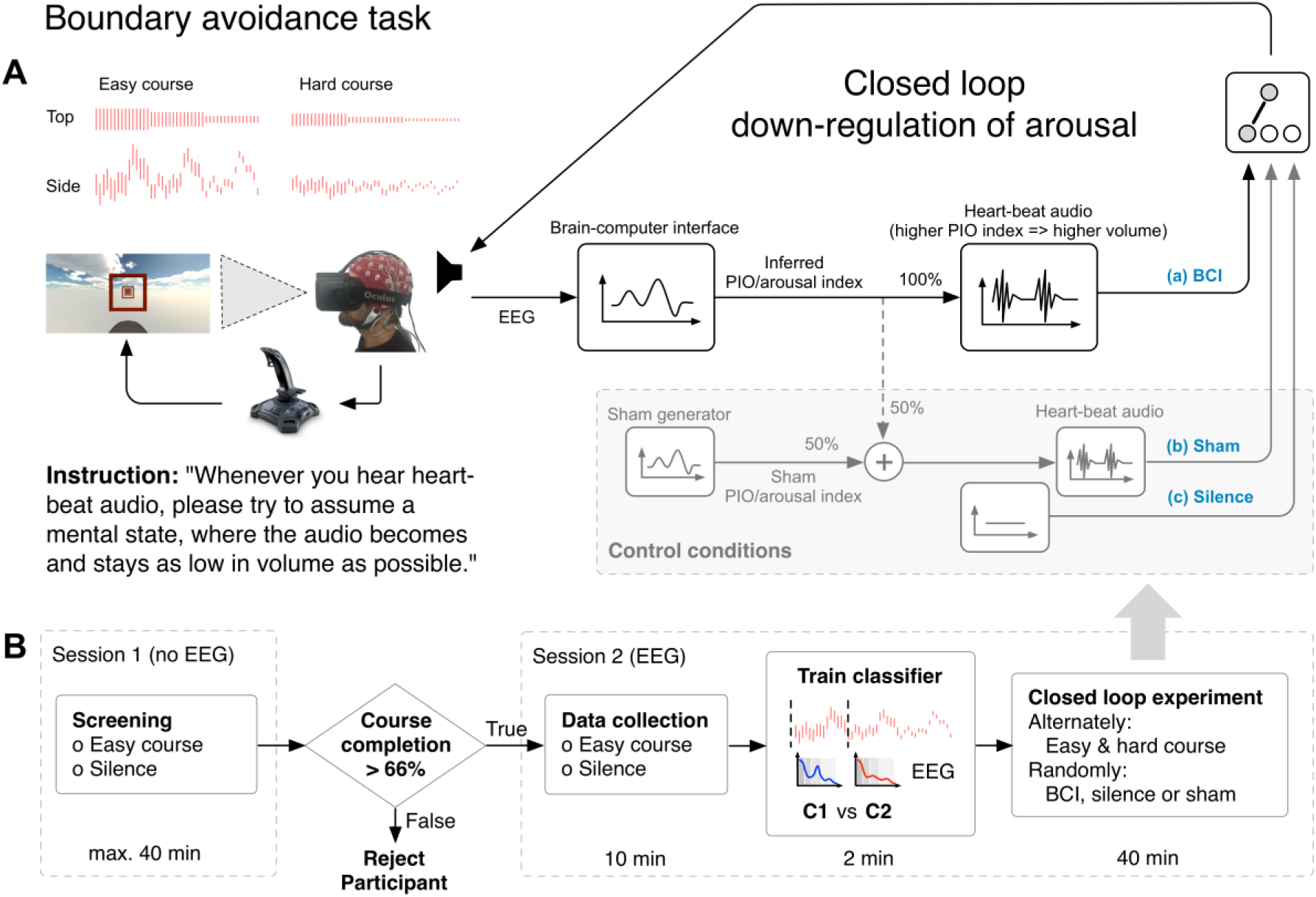
Setup of experiment and study protocol. (A) Study participants alternately guided a virtual aircraft through an easy or hard course of red rectangular boundaries (“rings”). Both courses were a maximum of 90 s long and increased in difficulty over time as ring sizes decreased. Missing a ring ended the flight trial immediately. Every new flight attempt was randomly assigned one of three feedback conditions: In the main condition (a) BCI (brain-computer interface), audio feedback from an EEG-based decoder was presented to the participant (“closed-loop” experiment). During the two control conditions (b) sham and (c) silence, partly random or no audio signal were presented respectively. Participants were instructed to down-regulate their arousal as outlined at the bottom left of the panel. (B) During initial screening in session 1, only novice participants able to repeatedly fly through 66 % of course type easy within 40 min were admitted for the main experiment in session 2. Session 2 started with 10 min of EEG collection while participants repeatedly attempted to fly through the easy course. The EEG-based decoder was then trained on this data, and subsequently used to generate feedback in the main EEG experiment.

### Neurofeedback improves flight performance under difficult conditions

In accordance with the Yerkes and Dodson law, we found BCI-based feedback to improve flight performance relative to control conditions for the unseen, untrained and more difficult flight course but not for the easier, previously trained course. This effect was reflected in a significant interaction between the independent variables *feedback* (levels: BCI, sham and silence) and *course* (levels: easy and hard) in an analysis of variance for the normalized dependent variable flight time (F_2,402_=3.535, P=0.031, R^2^=0.011). Post-hoc t-tests for course type hard, showed significantly prolonged normalized flight time for feedback type BCI relative to both control conditions silence and sham (Fig. 3A; descriptive statistics for raw flight time; tests on z-scored data; BCI: 46.2 ± 9.7 s (mean ± s.d.) vs silence: 39.0 ± 9.2 s; t_17_=-2.903, P=0.010, R^2^=0.331 and BCI vs sham: 38.2 ± 7.6 s; t_17_=-4.394, P<0.001, R^2^=0.532). Flight time with BCI feedback was thus on average increased by 18.3 % (i.e. 7.1 s) over silence where no feedback was provided and by 21.0 % (i.e. 8.0 s) over sham, where 50 % of the feedback signal were randomly generated and the other 50 % were true decoder output (Fig. 3B). No significant difference was found between the control conditions silence and sham in course type hard or between any of the three feedback conditions for course type easy (see Fig. 3C and 3D). These results are in agreement with our hypothesis, with the pattern of significance remaining unchanged even if corrected for multiple comparisons using a Holm-based correction (6 comparisons).

**Fig. 3.**
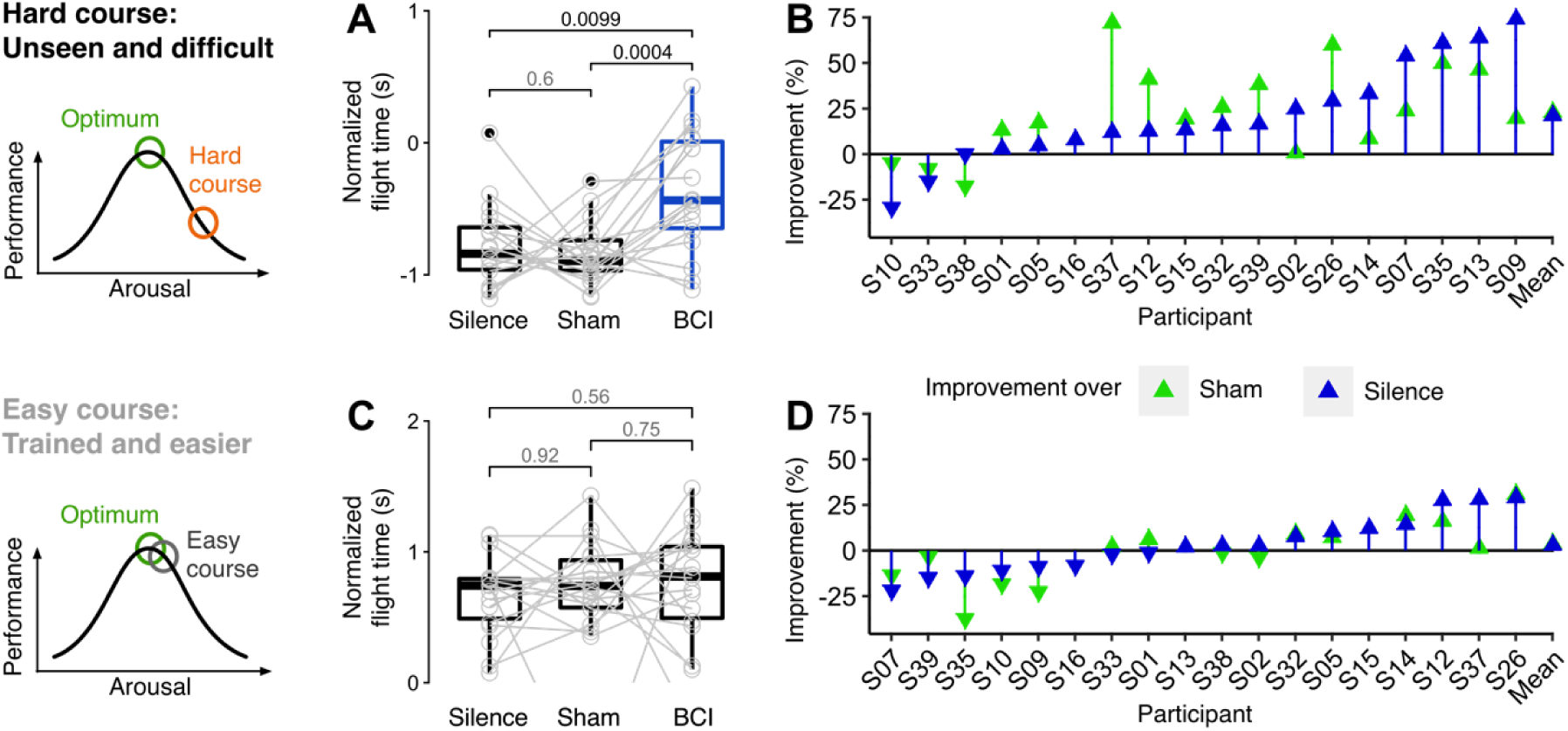
Flight performance improved with veridical neurofeedback in the hard but not the easy course. (A) Flight time increased significantly relative to both control conditions sham and silence for course type hard, where sharply elevated levels of arousal are expected. (B) In the hard course, individual flight performance consistently increased with veridical neurofeedback relative to controls for all except three subjects. (C) For the easy course, for which subjects were trained and screened and where no strongly elevated arousal was expected, no significant differences were found between any of the conditions. (D) Individual performance with veridical feedback relative to control conditions in course type easy showed no clear trend for a systematic increase or decrease. Hinges of boxplots represent first and third quartile and whiskers span from smallest to largest value of the data but reach out no further than 1.5 times the interquartile range. Numbers over brackets between boxplots represent uncorrected P-values of paired t-tests. The pattern of significance does not change after Holm-based correction for six comparisons.

The course-specific differences in performance improvement with BCI feedback could be explained by a difference in the baseline level of arousal for the two course types: Relative to the easy course, the unseen, untrained and more difficult course would hypothetically be associated with a higher level of arousal and consequently a higher potential for improvement via BCI-mediated down-regulation of arousal. We found evidence for higher task difficulty and arousal for course type hard relative to course type easy, manifested as significant differences in flight time (F_1,403_=366.529, P<0.001, R^2^=0.566), pupil size, heart rate and HRV (supplementary Fig. S2, S3 and S4). Consistent with our prediction based on the Yerkes-Dodson relationship, we observed improved flight performance relative to control conditions when subjects were instructed to down-regulate their arousal based on BCI feedback under difficult conditions.

### Higher heart rate variability for effective BCI feedback indicates lower arousal

In the hard course, we found significantly increased normalized HRV (metric pNN-35 ms) for BCI-based neurofeedback relative to the control conditions silence and sham (descriptive statistics for raw percent pNN-35 ms; tests on z-scored data; BCI: 41.8 ± 21.9 % (mean ± s.d.) vs silence 27.3 ± 19.4 %; t_13_=-4.514, P<0.001, R^2^=0.610; BCI vs sham: 30.6 ± 22.9 %; t_13_=-4.783, P<0.001, R^2^=0.638; Fig. 4A). No significant differences were found between the control conditions in course hard or between any of the conditions in course easy (Fig. 4C). The time domain based metric pNN-35 ms measures high frequency HRV (15). Increased high frequency HRV has been associated with increased activity of the parasympathetic nervous system along with decreased sympathetic nervous system activity and can be interpreted as decreasing arousal or stress (16). These results suggest that the observed improvement in task performance due to BCI feedback might be causally related to (or at least coincident with) a decrease in arousal experienced by the subjects.

**Fig. 4.**
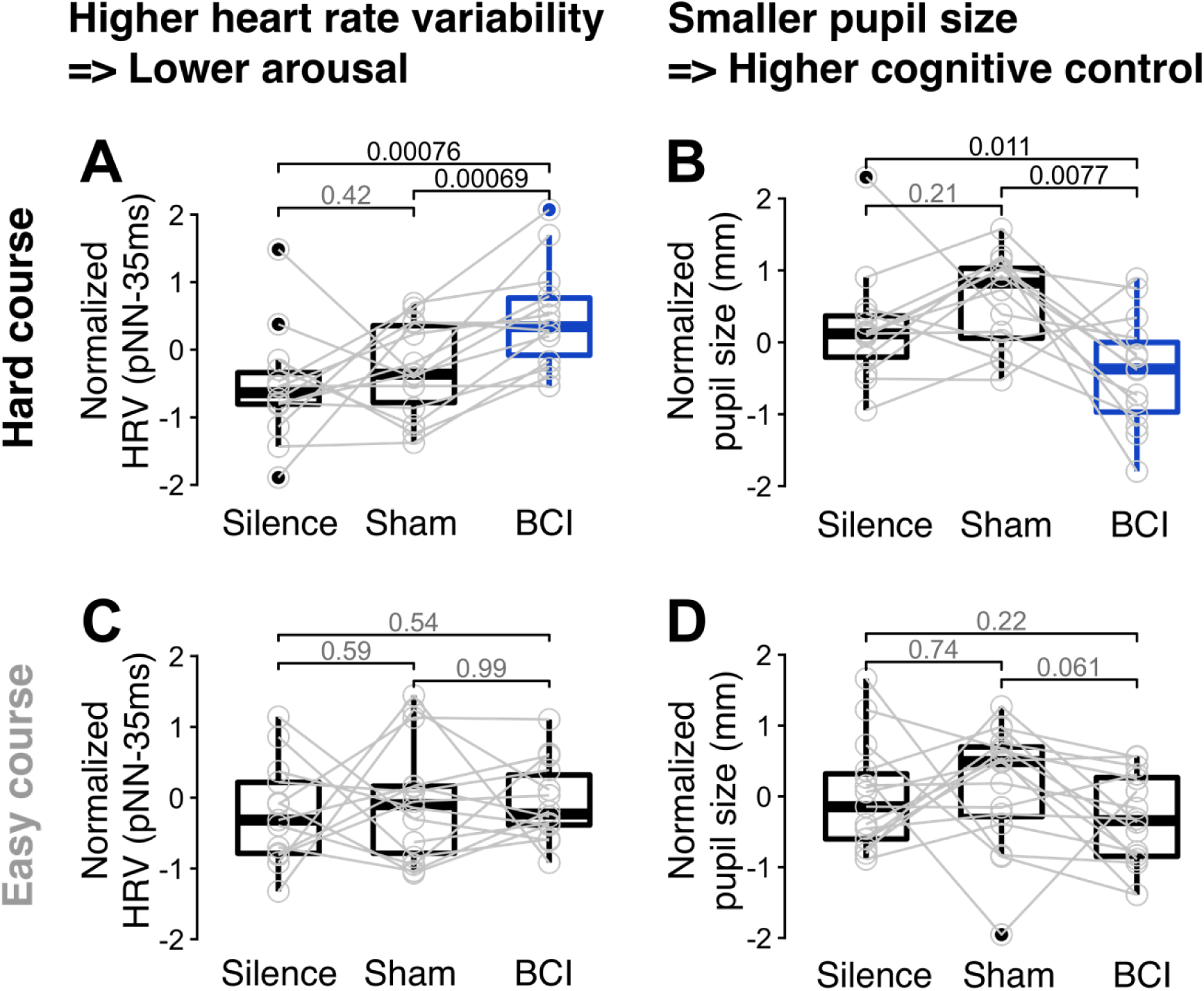
Significant changes in pupil size and heart rate variability (HRV) were observed in condition BCI relative to control conditions during course type hard. (A) For course type hard, HRV was significantly higher in condition BCI relative to both control conditions, while (B) normalized pupil size was significantly lower in condition BCI relative to both control conditions. For course type easy no significant effects were found for (C) HRV or (D) pupil size. Hinges of boxplots represent first and third quartile and whiskers span from smallest to largest value of the data but reach out no further than 1.5 times the interquartile range. Numbers over brackets between boxplots represent uncorrected P-values of paired t-tests. Holm-based correction for six comparisons does not change the pattern of significance for HRV but elevates the P-value of the paired comparison of normalized pupil size between conditions silence and BCI from P=0.011 to P_Holm_=0.057.

### Pupil activity implicates LC and ACC circuitry in performance improvement

Consistent with our prediction that lowered arousal would improve task performance by modulating LC activity, we found normalized pupil radius, a known correlate of LC activity, to be significantly decreased for the effective feedback condition in course type hard, relative to both control conditions silence and sham (Fig. 4B; descriptive statistic for raw radii; tests on z-scored data; BCI: 1.6 ± 0.2 mm (mean ± s.d.) vs silence: 1.7 ± 0.2 mm; t_12_=2.980, P=0.011, R^2^=0.425 and BCI vs sham: 1.7 ± 0.2 mm; t_12_=3.198, P=0.008, R^2^=0.460). No significant differences were found between the control conditions in course type hard or between any conditions in course type easy (Fig. 4D). Notably, with the rings in the paradigm being 2 s apart, these significant effects were only observed for the time window [1, 2] s after medium sized rings, but not for the 1 s time window directly following the rings (supplementary Fig. S6).

### Decoding task difficulty from the electroencephalogram

Decoding performance in cross validation on the training data set was 79.8 ± 7.2 % (mean ± s.d.; N=18) area under the receiver operator characteristics curve and statistically higher than chance for every participant (significance level, P=0.01; supplementary Fig. S8). Decoding performance for delta and theta band was significantly better than chance and when projecting the inverted linear decoder models onto the scalp (forward model) we observed a mid-frontal focus consistent with ACC involvement. Higher frequency bands like alpha, beta and gamma showed higher cross validated decoding performance and the associated forward models either reflected neural activity, non-neural activity (e.g. electromyogram) or a mix of both. This is consistent with previous findings (6) where neural activity was found in frontocentral delta and theta as well as parietal alpha. Similarly, Saproo and others also identified gamma activity that seemed to reflect changes in muscle tension. In accordance with the results from our preliminary study (N=3 subjects), cross-validated decoding performance was higher with EEG relative to joystick input.

## Discussion

We have shown that neurofeedback can be used to down-regulate arousal and improve human performance in a demanding sensory-motor task in real-time. This is in line with our prediction based on the Yerkes-Dodson law (11) and the theory of cognitive control (2). Cognitive control is a critical element of executive function that interacts with arousal systems to enable approach-avoidance behavior (18). When we avoid or “take flight” from a dangerous object, event and/or boundary, our avoidance response is reflexively in a direction opposite to that of the direction of where we perceive the danger — e.g. running away from the lion or the fire. The BAT scenario we study here is more complex in that avoidance of a boundary in one direction increases proximity to the boundary in another direction. The fundamental problem the brain is confronted with is one of a balancing act, where avoidance must be precisely controlled. The challenge is that this must occur in the presence of high-levels of arousal, which are believed to be forcing the brain to switch from exploiting previously learned models to stochastic exploration for new models (12, 14). Our findings in behavior and physiological signals support our hypothesis that the provided neurofeedback decreases arousal, by impeding this hypothetical switch to model exploration which we think then leads to the increased task performance.

Importantly, we found evidence in support of our hypothesis not only in increased task performance with veridical neurofeedback, but also in significant, concomitant changes in HRV and pupil size. Higher HRV with BCI feedback relative to control conditions indicates a lowering of arousal (16) and smaller pupil size under the same conditions can be interpreted as LC activity consistent with increased cognitive control (19). It is noteworthy, that significant effects in pupil size were only found in the epoch [1, 2] s after medium sized rings, but not in the epoch [0, 1] s. We interpret this pattern of pupil changes as reduced baseline pupil size and thus reduced tonic LC activity, along with increased task related pupil dilation and thus increased phasic LC activity that has been shown to follow correct task responses in monkeys (12). Overall, this pattern of pupil activity is consistent with a state of continued exploitation of previously learned models, high task engagement, high cognitive control and high task performance (12, 20). In accordance with the Yerkes-Dodson law, these effects were only present in the hard, unseen course where both arousal and task difficulty were expected to be higher than in the easy course. These findings are also supported by statistical analysis at a time resolution of 2 s (included in the supplemental material) showing that veridical feedback significantly predicted HRV, but only when subjects also heard the feedback and not in the silence condition. In other words, as subjects heard the feedback and attempted to down-regulate their arousal, successfully lowered feedback volume was associated with increased HRV, i.e. a lowering of arousal (15, 16). In summary, effects in HRV and pupil size, consistent with decreased arousal and a brain state of continued model exploitation were found only while veridical neurofeedback was provided in a difficult task and when task performance improved.

To our knowledge, this is the first system to improve human performance in a highly demanding sensory-motor task. Our system is novel in that it worked in real-time with a response time on the order of seconds. The system was otherwise unobtrusive and did not interfere with performance at moderate arousal levels. The only other study in literature where subjects concurrently modulated EEG-based feedback found increased performance in a sustained visual attention task under low levels of arousal lasting 120 minutes (21). A number of other studies reported that separate, dedicated neurofeedback training improved subsequent task performance. Using such approaches, fMRI-based studies showed improved performance in grip force control (22) or working memory (23) and EEG-based studies showed improved cognitive (24) or musical performance (25). A recent review on neurofeedback (26) points out that failures to replicate findings from promising preliminary studies in large clinical trials emphasizes the importance of understanding the physiological mechanisms that underlie neurofeedback approaches. We address this problem by complementing our report on a brain-behavior relationship, with supporting evidence across multiple conditions and physiological signals. Finally we contrast our approach against passive BCIs (27), which aim to improve human machine interaction by allowing a machine to unobtrusively adapt to covert aspects of user state like for example workload, surprise or fatigue. In one particularly interesting approach, the authors attempted to infer workload in eight subjects in real-time and activated machine assistance in a difficult visuomotor task whenever workload was high (28). This improved performance, but required the machine to know intricacies and successful control strategies for the task. Our approach requires no knowledge of task, environment or optimal control input but instead directly improves human performance. Given that passive BCIs require no active attention by the user, both approaches can be used concurrently.

There are several noteworthy limitations in the broad interpretation of our results due to specific choices in experimental design and the implemented closed-loop system. First is that our investigation relies on EEG and that subjects needed to be screened to meet a minimum task performance level so that enough EEG could be collected to train the decoder. A more complex experimental design, where task difficulty is additionally adjusted to the individual skill level of the recruit may have allowed us to admit additional subjects into this EEG study. After careful consideration during experimental design, we opted for the present approach since we thought it represented a good balance between complexity of setup and statistical analyses, logistic feasibility and experimental control. Another limitation of EEG is that its signal to noise ratio can decrease with increased environmental or biological noise such as muscle activity in the face or neck. One participant, changed their posture in the middle of the experiment, introducing so much muscle artifacts into the EEG that the data set became unusable and was excluded from analysis. For participant S10, decoder performance was unusually low at 66.7 % AUC compared to an average of 81.7 ± 7.2 % for all others. The subject was not excluded, but later presented with low performance in the BCI condition (Fig. 3B). Conceivably, degraded EEG signal quality could lower feedback efficacy. While there are promising approaches to improve EEG signal quality in the presence of noise (29), it is clear that using EEG in real-world applications could be challenging.

Another limitation is that even though we implemented our BAT paradigm in VR, it is still a simplified version of what one might expect in a real flight situation that would generate a PIO. For example, absent was simulated instrumentation that the pilot would direct their attention to when trending toward to a PIO. This increased attention toward the information in the instrumentation is thought to be a source of the increased cognitive workload (3–5) that generates a PIO (3–5). Our experiments assume that as flight difficulty increases during a trial, arousal levels increase and that this shifts the subject on the Yerkes-Dodson curve. We therefore do not directly address questions related to cognitive workload, rather our work is specific to arousal changes that couple to performance.

Our work also investigated only increases in arousal — i.e. the right side of the Yerkes-Dodson curve — and how arousal can be reduced to reach an operating point that optimizes task performance. The left side of the Yerkes-Dodson curve is also of practical interest, since it addresses cases of low arousal and fatigue that are also detrimental to task performance. Examples would include drowsiness, where one would want to increase arousal levels to improve task performance. Though our BAT experiments do not consider these low arousal states, we believe our approach potentially can be generalized to regulate arousal across the full range of the Yerkes-Dodson curve.

The collected evidence across conditions and multiple signal modalities furthered our understanding of the mechanisms underlying the efficacy of the presented approach, and suggests that this approach should generalize to other task domains where humans are forced to behave according to internal models of the environment under high arousal. Driving under difficult conditions, as another continuous visuomotor task, would be an obvious example. From a translational perspective it would be a great advantage if the presented feedback effect could be achieved based on non-neural signals like from joystick or steering wheel input alone, since such signals are easy to record in a real-world environment like in a car. For the present experiment, it seemed clear that it was best to base our decoder on EEG since a preliminary study (N=3) had shown that neither joystick nor EMG derived from face, neck or the right lower arm achieved higher decoding performance. In fact the decoding performance obtained based on facial and neck EMG was not statistically better than chance. Our results here confirm our previous observation, where decoding performance was higher based on EEG than when joystick input was used as the underlying signal. In terms of potential applications outside human machine interaction, the demonstrated approach of administering neurofeedback, while monitoring conformity with experimental hypotheses across conditions and multiple signal modalities, could serve as a model to enable targeted treatment in mental illness, where there is increasing evidence that cognitive and/or emotion regulation could lead to clinically significant improvement (30).

## Materials and Methods

### Subjects

We recruited 40 right handed, neurologically normal adults in New York City (age 26.2 ± 4.4 (mean ± s.d.) years; 23 female). Only subjects who reported normal hearing, normal vision or vision that was corrected to normal with contact lenses were included. We excluded volunteers who reported using medication that might influence the experiment. Participants were compensated with USD 20 per hour. After screening, 13 subjects were excluded, where for 12 subjects performance was too low, and one subject was non-novice to the task. Seven more could not enroll in the main session for other reasons (3 experienced VR sickness; for 2, technical problems prohibited recording; 1 subject’s hair-style made it impossible to record EEG of sufficient quality; 1 did not have time for the main experiment after all). Of 20 subjects (10 female) who were enrolled in the main study, 2 were excluded from analysis as one of them had to leave after less than half the session and the other changed their posture in the middle of the experiment, introducing so much muscle artifacts that the data set became unusable, leaving data from 18 subjects for analysis (age 24.9 ± 3.6; 8 female). Our experimental design used within-subject comparison, thus no randomization was used to assign subjects. The sequence of the three main conditions of interest silence, sham and BCI in the main experiment was random, but made sure that every condition occurred twice within six consecutive flight attempts. This study was approved by the institutional review board at Columbia University and written informed consent was obtained from all participants prior to screening and the main experimental sessions.

### Boundary avoidance task

Participants were instructed to fly a virtual plane through two different courses (*easy* and *hard*) of red boundaries (“rings” or “boxes"; Fig. 2A, top left). The vertical position of the rings was arranged along a trajectory computed as a sum of three sines. With the plane moving at a constant velocity and ~2 s of flight time between rings, both courses were a maximum of 90 s long while the size of the rings decreased every 30 s thus increasing difficulty. Consequently, courses consisted of three segments of large, medium and small rings. Flight attempts ended abruptly whenever a participant missed a single ring, after which the next flight attempt would be started. Course easy and hard each had a different, but fixed trajectory. Boundary sizes were overall smaller in the hard course rendering it more difficult. Participants controlled only the pitch of the virtual aircraft via right hand joystick input (A-10C HOTAS Warthog, Thrustmaster, France). Having a single degree of freedom (airplane pitch) was done to reduce the complexity of the experiment while still enabling a bit of realism. Joystick input was delayed by 0.2 s and non-linearly dampened to more closely resemble the more challenging flight characteristics of a real aircraft and consequently also increase the probability for PIOs to occur (6, 5). Virtual reality was used as the presentation mode since the looming of the glide boxes and their position was perceptually augmented by the immersiveness and binocular/stereo head mounted display, thus enabling strong modulation of arousal levels.

### Feedback conditions

Overall three different audio-based feedback conditions - silence, sham and BCI - were used throughout this study: In the first condition, *silence*, no audio was presented (i.e. Ω = 0 in Eq. (1)). The silent condition was important for both being an important control condition for the experiment (i.e. critical for showing that veridical neurofeedback improved performance over no neurofeedback at all) while also enabling additional analysis related to how the decoder output tracked other variables (HRV and pupil dilation) when subjects heard the decoder output.

The fixed combination of silence and the easy course were used during screening in session 1 and at the beginning of session 2 during 10 min of data collection while participants attempted flights. Based on these 10 min of data, a linear subject-specific decoder was trained to translate spontaneous EEG activity into an index of inferred task-dependent arousal between 0 and 100 %. For condition *BCI* this index of inferred arousal was temporally smoothed using a sliding window that was 5 s wide and instantaneously mapped onto the volume of low-rate (60 beats/min) synthetic heartbeat audio signal. That way, higher task difficulty and thus presumably higher task-dependent arousal corresponded to louder audio and vice versa (i.e. Ω = 1 and λ = 1 in Eq. (1)). For condition *sham*, the index of inferred task-dependent arousal was linearly combined with a randomly generated signal (AR_BCI; range also 0 to 100 %), such that Ω = 1 and λ = 0.5 in Eq. (1). More specifically, this random signal was generated as novelty observations from an autoregressive model (see Section Sham feedback). The average sound pressure levels for minimum and maximum loudness levels of feedback measured from the headphones were 59.6 dB and 71.1 dB respectively (measured via application Decibel X, 6.0 on iPhone 7, iOS 11.0.2). In the main part of the study, “Closed Loop Experiment” (Fig. 2B), participants alternately attempted flights in course easy and hard, while one of the three feedback conditions (BCI, sham or silence) was assigned randomly for every new flight attempt. Every feedback condition occurred twice in six flight attempts. In total, 24 flight attempts were recorded for every participant in the closed-loop block of the experiment. BCI was the main condition of interest, while sham and silence served as control conditions.

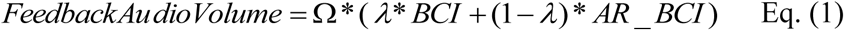

Our rationale for using the loudness of a low-rate heart beat as the mode for feedback was based on both the literature as well as pilot experiments evaluating other modes of feedback. The relationship between arousal, heart rate and the role of the locus coeruleus (LC) in regulating both cortical arousal and parasympathetic neurons that control heart rate has been established (31) and thus pointed us to using low-rate heart beat as a mode of feedback to present to the subject. We also conducted pilot experiments to evaluate other modes for presenting the neurofeedback. Specifically we tested visual presentation of neurofeedback via a vertical temperature gauge that changed in height and color (green to red) as the neurofeedback output increases. We also tested a mode of feedback whereby we adaptively adjusted the control/response parameters of the joystick, via software, such that high neurofeedback reduced the gain between the stick movement and movement of the simulated aircraft. For both the visual feedback and the control-based feedback, subjects performed substantially worse with neurofeedback than in the silent case. Given the literature and these pilot experiments, we chose the auditory feedback of low-rate heart beats for our experiments.

### Instruction

Participants were instructed based on slides and further instructions were read by the experimenter before a new experimental block was started. We explained, that the purpose of this study was to investigate brain activity that was elicited by the BAT paradigm and that audio feedback was provided based on the subject’s current brain activity. We kept participants blind to our aim of investigating flight performance differences and the existence of the condition sham. The key instructions were as follows: “Go through every box!”, “Consider missing a box the equivalent of crashing a plane” and “Whenever you hear heartbeat audio, please try to assume a mental state where the audio becomes and stays as low in volume as possible”.

### Screening

Prior to the main EEG experiment, all 40 recruits were admitted to a training and screening session where no EEG was recorded. This had two reasons: First, to assure a comparable baseline level of task proficiency across subjects and second, to make sure subjects were able to fly far enough through the easy course so that later in the main experiment, enough EEG could be collected to calibrate the BCI. For a maximum of 40 minutes, subjects were allowed to make flight attempts in the easy course, while no feedback was provided (condition silence). Subjects wore headphones with noise-canceling activated and we recorded joystick input and pupil diameter during flight attempts. The threshold criterion for passing screening was met by completing or exceeding 66 % of the 90-second course in 3 out of 4 consecutive attempts. One subject was found to be non-novice to the task and was thus excluded.

### Decoder for real-time feedback

Based on EEG collected during 10 min of flight attempts in course easy at the beginning of session 2, we trained a subject specific, multivariate linear model that indexed inferred task-dependent arousal between 0 and 100 %, so that a high index value corresponded to high task dependent arousal (thus presumably also low cognitive control) and vice versa. The setup was initially treated as binary classification problem: Class 1 was represented by EEG data collected during the first segment of the flight course where boundary size was largest and task difficulty was lowest. Class 2 was represented by EEG data collected during the second and third segment of the course, where boundary sizes were smaller and task difficulty was consequently higher. Every 2 s epoch of EEG between rings was treated as a separate observation. The linear decoding model was then obtained in two steps, where first, linear projections from EEG space into a six dimensional surrogate subspace were computed via filter bank common spatial patterns (FBCSP (32)). These subspace projections were computed separately for every one of the five frequency bands 0.5 to 4, 4 to 8, 8 to 15, 15 to 24, 24 to 50 Hz, such that variance-based between-class separability was maximized. The subspace projections were attained by first computing eigendecomposition of *M_c_* in Eq. (2) separately for class 1 (c = 1) and class 2 (c = 2)

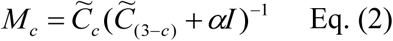

where 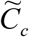 was the channel covariance matrix (size 64×64) for class c, *α* = 10^−10^ was a Tikhonov regularization parameter (33) obtained empirically based on previously collected data (6) and was the identity matrix (size 64×64). For every band, and for every class c = {1, 2}, only the three eigenvectors of *M_c_* that were associated with the largest eigenvalues were retained. Thus a projection matrix of size 64 × 6 was obtained for every frequency band which overall resulted in a 30 dimensional feature space (64 EEG channels; 6 eigenvectors per band × 5 frequency-bands → 30 features). In step two, this 30 dimensional feature space after FBCSP processing was projected down to a scalar dimension using shrinkage regularized linear discriminant analysis (LDA (34)). A scaling parameter was obtained by normalizing the LDA output for all training data and stored with the model so that real-time output could be scaled accordingly. The decoder output was subjected to temporal smoothing using a window that was 5 seconds wide to reinforce neural activity and to suppress noise and spurious fluctuations.

### Sham feedback

To generate sham feedback in real-time, the feedback which was usually only based on EEG (BCI) was linearly combined with simulated novelty observations from an autoregressive (AR) model (c.f. 35, Eq. (3); signal AR_BCI in Eq. (2)). The AR model had been trained based on nine data sets from a previous study where EEG was recorded while study participants attempted to fly through the same two courses as in this study but in the absence of closed loop feedback (6). For AR model setup, FBCSP had first been trained for every single subject on all the EEG data of every single subject. Subsequently the individual FBCSP decoder models were applied to the EEG data of the same subject to obtain subject specific time series of an index of inferred task-dependent arousal at a sampling rate of 16 Hz. Then, AR models of orders 5 to 80 (in steps of 5) were fit separately for every subject’s time series based on Burg’s method (36). AR model order p = 40 yielded the lowest average Bayesian Information Criterion score across participants and was thus selected for the final model. The coefficients for the final AR model in Eq. (3), φ_1_ … φ_p_ along with constant offset c were determined by averaging coefficients and offset of subject-specific models of order p = 40 across participants. For real-time generation of AR model based sham signal, the model was initialized with zeros and a random noise term ε_t_, drawn from a Gaussian distribution, was added for every prediction. No initialization effects were apparent latest after 30 s but the sham generator typically ran more than 5 min before its signal was provided as part of sham feedback.

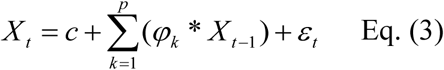

### Setup and signal acquisition

Subjects were seated comfortably inside a Faraday cage, wearing a head-mounted virtual reality display (Oculus Rift DK2, Oculus VR LLC, Menlo Park, CA, USA) and noise cancelling headphones (QuietComfort 20, Bose Corporation, MA, USA). The 3D paradigm was designed using NEDE (37), a scripting framework to design experiments in virtual 3D environments based on Unity (Unity Technologies, San Francisco, CA, USA). We used the software lab streaming layer (38) to synchronize acquisition of signals of different sampling rates (f_s_). In session 1 we recorded joystick input (f_s_=60 Hz), pupil radius and eye gaze from an eye-tracker within the head-mounted headset (SensoMotoric Instruments, Germany; f_s_=60 Hz), paradigm markers and flight trajectories (f_s_=75 Hz). In session 2, we additionally recorded 64 channels of EEG, the electrocardiogram from two electrodes placed on the thorax, electrodermal activity from two electrodes placed on the inside of the left hand and respiration from a belt around the thorax (ActiveTwo biosignal amplifier, BioSemi B.V., Netherlands; f_s_=2048 Hz).

### Statistical analysis

The dependent variable flight time was normalized to zero mean and unit variance within each subject to account for individual differences, before a linear model was fitted using ordinary least squares regression in R (Version 3.3.3 (39)) using categorical and continuous predictors including course difficulty (easy and hard), feedback condition (silence, sham and BCI), and the interaction of the two. Further predictors were subject demographics including age, gender, hours slept the previous night, average weekly gaming hours over the last three years, number of screening trials, and subject-derived continuous signals related to power of the joystick input signal, heart rate, BCI output and pupil size. To satisfy the requirement for normality of the residuals, we iteratively removed outliers by visual inspection of diagnostic plots in R, including scatter plots of fits vs. residuals, QQ plots, leverages vs. residual plots relative to Cook’s distance, and scale location plots, until the residuals met statistically tested requirements (R package GVLMA, version 1.0.0.2, (40)). Subsequently, the previously fitted linear model was subjected to analysis of variance. Post-hoc tests for the difference of means were computed using paired, two-sided Student’s t-tests, where equal variance was not assumed. Flight length within subject was collapsed for post-hoc tests based on the median. Pupil size and HRV were extracted from these exact flight attempts of median length. Between-course differences were preserved in the normalization of flight time, enabling analysis of the interaction of course and feedback condition. Pupil and HRV were additionally normalized within-course as our main interest was in comparing these physiological signals between feedback conditions. Regardless of which type of normalization is used, the patterns of significance remain unchanged. If not stated otherwise, descriptive statistics are reported as mean ± standard deviation. Statistical effects were considered significant for P<0.05. We encourage to interpret uncorrected P-values of post-hoc tests relative to our specific hypotheses, but also report results of Holm based correction for reference. Sample size was determined a priori to be 20 participants assuming partial η^2^=0.5, α=0.05 and 1-β=0.95 for the two main effects course and condition and their interaction (software G*Power 3.1.9.2 (41), University of Düsseldorf, Germany).

## Acknowledgments

### General

The authors thank Yida Lin for help with figures and formatting and James McIntosh for helpful discussion.

### Funding

The work was funded by the Defense Advanced Research Projects Agency (DARPA) and Army Research Office (ARO) under grant number W911NF-11-1-0219, the Army Research Laboratory under Cooperative agreement number W911NF-10-2-0022, the National Science Foundation under grant IIS-1527747 and the UK Economic and Social Research Council under grant number ES/L012995/1. The views and conclusions contained in this document are those of the authors and should not be interpreted as representing the official policies, either expressed or implied, of the US Government. The US Government is authorized to reproduce and distribute reprints for Government purposes notwithstanding any copyright notation herein.

### Author contributions

J.F., P.S. and S.S. designed the experiment. J.F. implemented the experimental setup. J.F. and J.C. collected the data. J.F. analyzed the data. J.F. and P.S. wrote the paper. All authors contributed to the editing of the final manuscript.

### Competing interests

The authors declare that they have no competing financial interests.

### Data and materials availability

All data is available on reasonable request.

## References

1. Sallet, J., Mars, R., Quilodran, R., Procyk, E., Petrides, M., & Rushworth, M., Neuroanatomical basis of motivational and cognitive control: a focus on the medial and lateral prefrontal cortex. *In* Mars, R. (ed.), Neural Basis of Motivational and Cognitive Control (MIT Press, Cambridge, MA, 2011).

2. Botvinick, M. M., Cohen, J. D. & Carter, C. S., Conflict monitoring and anterior cingulate cortex: an update. Trends in Cognitive Sciences 8, 539–546 (2004).

3. Gray, W. R., Boundary-avoidance tracking: a new pilot tracking model. In AIAA Atmospheric Flight Mechanics Conference and Exhibit, San Francisco, California, 86–97 (2005).

4. Dotter, J. D., An Analysis of Aircraft Handling Quality Data Obtained From Boundary Avoidance Tracking Flight Test Techniques. Master’s thesis (2007).

5. Gray, W. R., A generalized handling qualities flight test technique utilizing boundary avoidance tracking. In US Air Force T&E Days, Los Angeles, 1648 (2008).

6. Saproo, S., Shih, V., Jangraw, D. C. & Sajda, P., Neural mechanisms underlying catastrophic failure in human-machine interaction during aerial navigation. Journal of Neural Engineering 46, 133–143 (2016).

7. Botvinick, M. M., Braver, T. S., Barch, D. M., Carter, C. S. & Cohen, J. D., Conflict monitoring and cognitive control. Psychological review 108, 624–652 (2001).

8. Cavanagh, J. F. & Frank, M. J., Frontal theta as a mechanism for cognitive control. Trends in Cognitive Sciences 18, 414–421 (2014).

9. Foxe, J. J. & Snyder, A. C., The Role of Alpha-Band Brain Oscillations as a Sensory Suppression Mechanism during Selective Attention. Frontiers in psychology 2 (2011).

10. Green, J. J., Boehler, C. N., Roberts, K. C., Chen, L. C., Krebs, R. M., Song, A. W., & Woldorff, M. G., Cortical and subcortical coordination of visual spatial attention revealed by simultaneous EEG-fMRI recording. The Journal of neuroscience 37, 7803–7810 (2017).

11. Yerkes, R. M. & Dodson, J. D., The relation of strength of stimulus to rapidity of habit-formation. Journal of comparative neurology and psychology 18, 459–482 (1908).

12. Aston-Jones, G. & Cohen, J. D., An integrative theory of locus coeruleus-norepinephrine function: adaptive gain and optimal performance. Annu. Rev. Neuroscience 28, 403–450 (2005).

13. Wolpaw, J. R. & Wolpaw, E. W. Brain-Computer Interfaces: Principles and practice (Oxford University Press, 2012).

14. Tervo, D. G, Proskurin, M., Manakov, M., Kabra, M., Vollmer, A., Branson, K., & Karpova, A. Y., Behavioral variability through stochastic choice and its gating by anterior cingulate cortex. Cell 159, 21–32 (2014).

15. Task Force of the European Society of Cardiology, Heart rate variability: standards of measurement, physiological interpretation and clinical use. Circulation 93, 1043–1065 (1996).

16. Berntson, G. G. & Cacioppo, J. T., Heart rate variability: Stress and psychiatric conditions. *In* Malik, M. & Camm, A. J. (eds.), Dynamic electrocardiography, 57–64 (John Wiley & Sons, 2004).

17. Pfurtscheller, G., Allison, B. Z., Brunner, C, Bauernfeind, G., Solis-Escalante, T., Scherer, R., Zander, T. O., Müller-Putz, G., Neuper, C. & Birbaumer, N., The hybrid BCI. Frontiers in Neuroscience 4, 30 (2010).

18. Koch, S., Holland, R. W. & van Knippenberg, A., Regulating cognitive control through approach-avoidance motor actions. Cognition 109, 133–142 (2008).

19. Joshi, S., Li, Y., Kalwani, R. M. & Gold, J. I., Relationships between pupil diameter and neuronal activity in the locus coeruleus, colliculi and cingulate cortex. Neuron 89, 221–234 (2016).

20. Gilzenrat, M. S., Nieuwenhuis, S., Jepma, M. & Cohen, J. D., Pupil diameter tracks changes in control state predicted by the adaptive gain theory of locus coeruleus function. Cognitive, Affective and Behavioral Neuroscience 10, 252–269 (2010).

21. Beatty, J., Greenberg, A., Deibler, W. P. & O’Hanlon, J. F., Operant Control of Occipital Theta Rhythm Affects Performance in a Radar Monitoring Task. Science 183, 871–873 (1974).

22. Blefari, M. L., Sulzer, J., Hepp-Reymond, M. C., Kollias, S., & Gassert, R., Improvement in precision grip force control with self-modulation of primary motor cortex during motor imagery. Frontiers in Behavioral Neuroscience 9 (2015).

23. Sherwood, M. S., Kane, J. H., Weisend, M. P. & Parker, J. G., Enhanced control of dorsolateral prefrontal cortex neurophysiology with real-time functional magnetic resonance imaging (rt-fMRI) neurofeedback training and working memory practice. Neuroimage 124, 214–223 (2016).

24. Hanslmayr, S., Sauseng, P., Doppelmayr, M., Schabus, M. & Klimesch, W., Increasing individual upper alpha power by neurofeedback improves cognitive performance in human subjects. Applied Psychophysiology and Biofeedback 30, 1–10, (2005).

25. Egner, T. & Gruzelier, J. H., Ecological validity of neurofeedback: modulation of slow wave EEG enhances musical performance. Neuroreport 14, 1221–1224 (2003).

26. Sitaram, R., Ros, T., Stoeckel, L., Haller, S., Scharnowski, F., Lewis-Peacock, J., Weiskopf, N., Blefari, M. L., Rana, M., Oblak, E. & Birbaumer, M., Closed-loop brain training: the science of neurofeedback. Nature Reviews Neuroscience, 86–100 (2017).

27. Zander, T. O. & Kothe, C. Towards passive brain-computer interfaces: applying brain-computer interface technology to human-machine systems in general. Journal of Neural Engineering 8, 025005 (2011).

28. George, L., Marchal, M., Glondu, L. & Lécuyer, A., Combining brain-computer interfaces and haptics: detecting mental workload to adapt haptic assistance. In International conference on human haptic sensing and touch enabled computer applications, 124–135 (2012).

29. Daly, I., Scherer, R., Billinger, M. & Müller-Putz, G., FORCe: Fully online and automated artifact removal for brain-computer interfacing. IEEE Transactions on Neural Systems and Rehabilitation Engineering 23, 725–736 (2015).

30. Keynan, J. N., Meir-Hasson, Y., Gilam, G., Cohen, A., Jackont, G., Kinreich, S., Ikar, L., Or-Borichev, A., Etkin, A., Gyurak, A. & Klovatch, I., Limbic activity modulation guided by fMRI-inspired EEG improves implicit emotion regulation. Biological Psychiatry 80, 490–496 (2016).

31. Wang, X., Pinol, R. A., Byrne, P., Mendelowitz, D., Optogenetic Stimulation of Locus Ceruleus Neurons Augments Inhibitory Transmission to Parasympathetic Cardiac Vagal Neurons via Activation of Brainstem alpha 1 and beta 1 Receptors. Journal of Neuroscience 34, 6182–6189 (2014).

32. Ang, K. K., Chin, Z. Y., Zhang, H. & Guan, C., Filter Bank Common Spatial Pattern (FBCSP) in brain-computer interface. In IEEE International Joint Conference on Neural Networks, Hong Kong, 2390–2397 (2008).

33. Lotte, F. & Guan, C., Regularizing common spatial patterns to improve BCI designs: unified theory and new algorithms. IEEE Transactions on Biomedical Engineering 58, 355–362 (2011).

34. Ledoit, O. & Wolf, M., A well-conditioned estimator for large-dimensional covariance matrices. Journal of Multivariate Analysis 88, 365–411 (2004).

35. Hamilton, J. D., Time Series Analysis (Princeton University Press, 1994).

36. Burg, J. P., Maximum entropy spectral analysis. In 37th Annual international meeting of the Society for Exploratory Geophysics, Oklahoma City (1967).

37. Jangraw, D. C., Johri, A., Gribetz, M. & Sajda, P., NEDE: An open-source scripting suite for developing experiments in 3D virtual environments. Journal of neuroscience methods 235, 245–251 (2014).

38. Kothe, C. A., Labstreaming layer (LSL): A system for unified collection of measurement time series in research experiments (2013). URL: https://github.com/sccn/labstreaminglayer/.

39. R Core Team. R: A Language and Environment for Statistical Computing. R Foundation for Statistical Computing, Vienna, Austria (2013). URL: http://www.R-project.org/.

40. Peña, E. & Slate, E., Global validation of linear model assumptions. Journal of the American Statistical Association 101, 341–354 (2006).

41. Faul, F., Erdfelder, E., Lang, A.-G. & Buchner, A., G*power 3: A flexible statistical power analysis program for social, behavioral, and biomedical sciences. Behavior Research Methods 39 (2007).

42. Billinger, M., Daly, I., Kaiser, V., Jin, J., Allison, B. Z., Müller-Putz, G. R., & Brunner, C., Is It Significant? Guidelines for Reporting BCI Performance. In Allison, B. Z., Dunne, S., Leeb, R., Del R. Millan, J. & Nijholt, A. (eds.) Towards Practical Brain-Computer Interfaces, 333–354 (Springer Berlin Heidelberg, 2013).

